# *Mitf* over-expression leads to microphthalmia and coloboma in *Mitf-cre* mice

**DOI:** 10.1101/2024.07.24.605021

**Authors:** Anne Nathalie Longakit, Hannah Bourget, Catherine D. Van Raamsdonk

## Abstract

The Microphthalmia associated transcription factor (Mitf) is a critical regulator of the melanocyte lineage and also plays an important role in eye development. Mitf activity in different cell types is controlled in part by ten alternative promoters and their resulting isoforms. A useful tool for melanocyte-based research, the *Mitf-cre* transgene was designed to express Cre recombinase from the Mitf-M promoter, which is melanocyte specific. However, *Mitf-cre* mice are also microphthalmic, perhaps because of insertional mutagenesis or disrupted gene expression. Here, we investigated these possibilities. We determined that the eye phenotype arises early, with *Mitf-cre* embryos at E13.5 exhibiting variable ocular sizes and abnormalities, but all with coloboma. Targeted locus amplification and next generation sequencing indicated that multiple copies of the transgene integrated into an intergenic region on chromosome 2, in between *Spred1* and *Meis2*. The BAC transgene used to make *Mitf-cre* was larger than expected, carrying three upstream alternative promoters, Mitf-H, Mitf-D, and Mitf-B, which could express their isoforms intact off the transgene. RT-qPCR using eye tissue demonstrated a 5-fold increase in *Mitf* transcripts containing exon 1B1b, which is shared by Mitf-H, Mitf-D, and Mitf-B, while *Spred1* and *Meis2* did not differ in their expression. These findings clarify and support the usage of *Mitf-cre* in conditional mutagenesis in melanocytes. The specific over-expression of the Mitf-H and Mitf-D isoforms, which are preferentially expressed in the RPE, presents a unique resource for those interested in eye development and coloboma.

## INTRODUCTION

The microphthalmia associated transcription factor (*Mitf*) was first described in 1993 as a basic helix-loop-helix leucine zipper protein by Hodgkinson *et al*^1^. The authors determined that this gene was responsible for the classic mouse mutant, *Microphthalmia* (*Mi*), which includes pigmentary defects, small eyes (microphthalmia), osteopetrosis, deafness, and mast cell deficiency ^1–4^. Indeed, mutations in *MITF* produce similar manifestations in humans, resulting in the Waardenburg and Tietz syndromes^5,6^. In *Mitf*-null mice, there is an absence of neural crest derived melanocytes and a failure of the retinal pigment epithelium (RPE) to differentiate^7–9^. Hence, Mitf is critical for both types of pigment producing cells found in the eye, melanocytes and the RPE.

Within the developing eye, Mitf works in concert with other transcription factors, such as Vsx2, Pax2, Tbx5, and Sox2, as well as eye field transcription factors, Pax6, Rax, Six6, Six3, and Lhx2, to generate well-defined expression domains during a series of morphogenic events that establish patterning across three ocular axes (dorso-ventral, distal-proximal, and nasal-temporal) (**Figure 1A**)^10–12^. In mice, this series of stereotypic events begins at embryonic day (E) 8.0 with the establishment of the eye field in the presumptive forebrain^10,13^. At E8.5, the diencephalon evaginates to form bilateral optic vesicles (OV). *Mitf* is expressed in the OV at this stage (**Figure 1B**)^10,14^. As the OV displaces the surrounding periocular mesenchyme and contacts the FGF-expressing overlying surface ectoderm, *Vsx2* (also called *Chx10*) is activated, which subsequently down-regulates *Mitf* in the distal-most OV^10,15–17^. This part of the OV will become the neural retina, after invagination along the optic fissure forms a two layered structure. The outer layer continues to express *Mitf* and forms the RPE ^10,12,13,18–20^.

**Figure 1.**
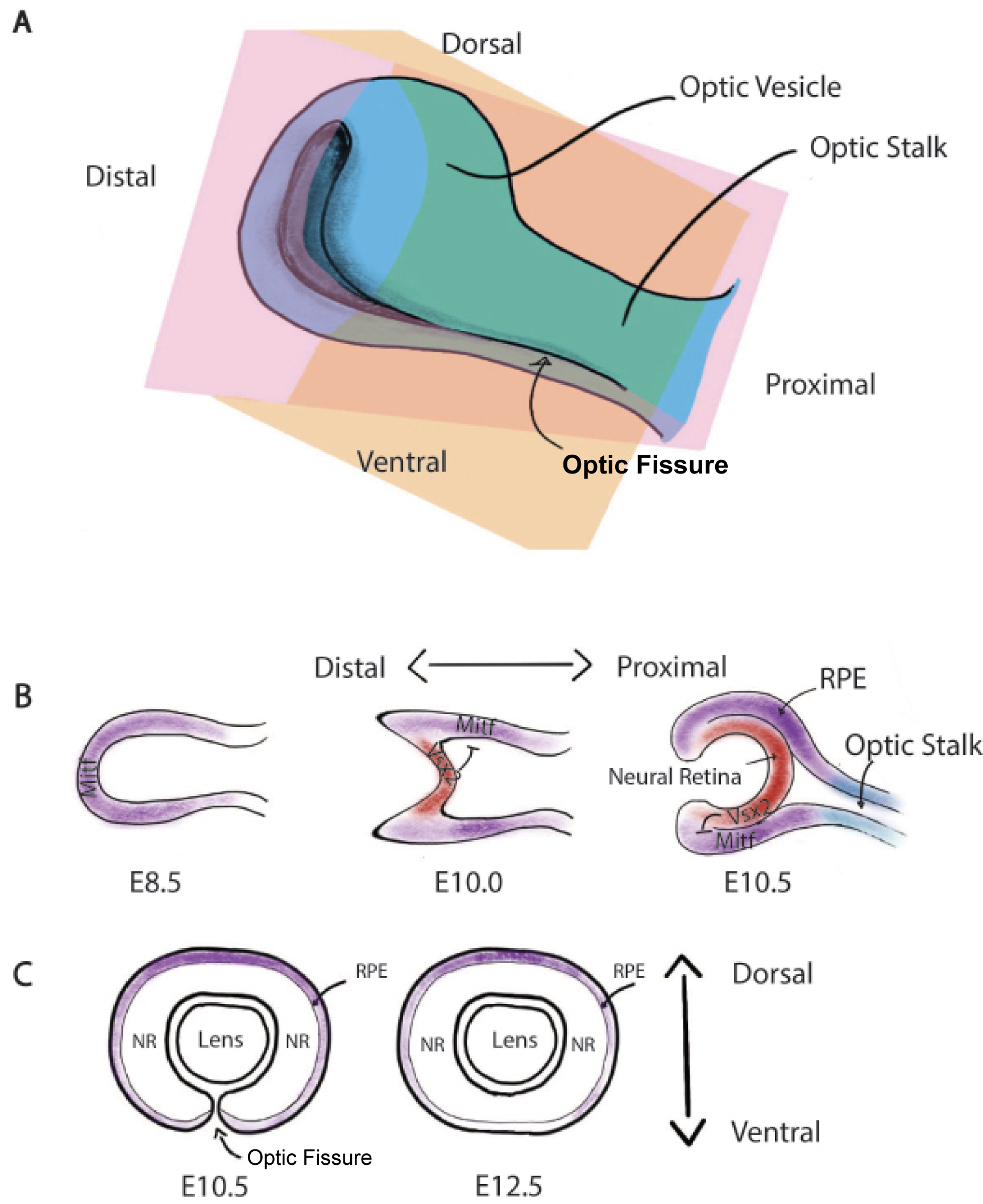
The expression of *Mitf* in ocular development. **(A)** Schematic of mouse ocular development at E10.5 depicting the coronal and sagittal planes of reference as orange and pink, respectively. **(B)** Mitf expression (purple) during mouse ocular development from E8.5-10.5, as viewed from the sagittal plane. Vsx2 (red) inhibits Mitf expression in the future neural retina. The optic stalk is shown in blue. **(C)** A gradient of Mitf expression (purple) during optic fissure closure at E10.5-12.5, as viewed from the coronal plane.

Meanwhile, constriction of the most proximal OV generates the prospective optic stalk, which expresses *Pax2*, again creating an expression boundary ^10,13,18^. On the ventral side of the OV, the optic fissure runs along the proximal-distal axis^12,13,19,21,22^. This fissure allows the necessary vasculature to enter and support the eye during development^15,17,22,23^. With further growth, the margins of the optic fissure become apposed and ultimately merge to complete optic fissure closure by E12.5 (**Figure 1C**)^22,24^. Subsequently, other tissues of the eye develop from the surrounding mesenchyme and melanocytes from the neural crest enter to populate the uveal tract. When fully developed, the eye is comprised of concentrically arranged tissue layers derived from different embryonic origins, with Mitf expression restricted to the RPE and the melanocytes^9,10,14^.

Due to the critical role of Mitf in ocular development, mutations in *Mitf* lead to a spectrum of structural defects such as microphthalmia, anophthalmia, and coloboma^11,19^. Horsford *et al.*^20^ found that *Mitf* deficiency in OV cultures generates ectopic neural retina tissue, whereas the ectopic expression of *Mitf* shifted the fate of OV cells from a neural retina phenotype to an RPE phenotype^20^. Furthermore, the failure to exclude *Mitf* from the margins of the optic fissure led to upregulated RPE fate and failed optic fissure closure, thus highlighting the importance of maintaining specific *Mitf* expression levels and boundaries during ocular development ^24^.

To add to this specificity, *Mitf* is regulated in an isoform selective manner, with preferential isoform expression occurring in specific tissue and cell types^14^. In humans and mice, there are ten Mitf isoforms with different N-termini, each of which is encoded by a separate first exon. The isoforms all share exon 1B1b, except for Mitf-M, which is the furthest downstream. All transcripts then splice to exon 2^14^. The Mitf-M isoform is exclusively expressed in melanocytes, but the other isoforms have more complicated expression patterns. In terms of the RPE, Mitf-A, Mitf-J, Mitf-H and Mitf-D are all expressed to varying degrees ^2,8,14^. Mitf-B has not been studied as much, but its start site is close to Mitf-H and Mitf-D. A deletion in Mitf-M exon 1 produced a loss of melanocytes, including those in the choroid, while maintaining pigmentation in the RPE^25^. Furthermore, examination of *Mitf ^mi-rw/mi-rw^*mice, which carry an 86 kb deletion covering the Mitf-H, Mitf-D, and Mitf-B promoter group, revealed dorsal RPE thickening, which subsequently transdifferentiated into a second neural retina^14^.

In 2007, Alizadeh *et al.* generated *Tg(Mitf-cre)7114Gsb (i.e.* "*Mitf-Cre")* mice, a transgenic line that expresses Cre recombinase under the control of the Mitf-M promoter as a tool to study pigment cell biology^26^. In order to include as much of the endogeneous gene expression regulatory sequences as possible, a mouse bacterial artificial chromosome (BAC) from chromosome 6 containing part of the *Mitf* locus was used as the vector (RP23-271N22). The *Mitf-Cre* transgene is especially useful with regard to its specificity and sensitivity. Indeed, evaluation of X-gal staining in E12.5 whole embryos and adult tail epidermis of progeny from a cross between the *Mitf-Cre* founder and mice carrying the Gt(Rosa)26^tm1Sor^ reporter gene (R26R), labeled well characterized melanoblast populations in the trunk paraspinal region and head. However, 100% of the *Mitf-cre* animals on the original C57Bl/6 genetic background had microphthalmia, which was unexpected. The cause of the eye phenotype was proposed to be transgene associated, but was not investigated^26^.

Subsequently, we backcrossed *Mitf-cre* to a different genetic background, C3HeB/FeJ, and found a significant decrease in the severity of the eye phenotype, which allowed the transgene to be used to study ocular melanoma driven by GNAQ^Q209L^ ^27^. In addition, we found that *Mitf-cre* was a potent driver for cutaneous melanoma driven by Braf^V600E^ ^28^. Because of its utility, it is important to understand the cause of the *Mitf-cre* eye phenotype; specifically whether it is something about the transgene itself or a disruption caused by the transgene’s random insertion into the genome. In addition, the microphthalmia phenotype may be interesting with respect to eye development. In this paper, we better characterized the eye phenotype of *Mitf-cre* on the C3HeB/FeJ background, mapped the location of the *Mitf-cre* transgene insertion into the mouse genome, and measured the expression of key genes of interest in the *Mitf-cre* eye.

## RESULTS

The original *Mitf-Cre* mouse line that was produced on the C57Bl/6 genetic background exhibited severe microphthalmia (closed eyelids) in all mice. Subsequent breeding in our lab involving the backcrossing and maintenance of the *Mitf-cre* line on the C3HeB/FeJ background resulted in a reduction in the severity of the eye phenotype, to open eyes or even normal sized eyes (**Figure 2A**)^27^. Some *Mitf-cre* mice had non-bilateral phenotypes, with one open and one closed eye ("mixed").This phenotypic difference may be due to genetic background specific effects. For example, previous reports have indicated that mice carrying a heterozygous *Pax6* mutation on the C57Bl/6 background had more severe microphthalmia compared to a hybrid genetic background, purportedly due to epistatic interactions between genotype and background^29^.

**Figure 2.**
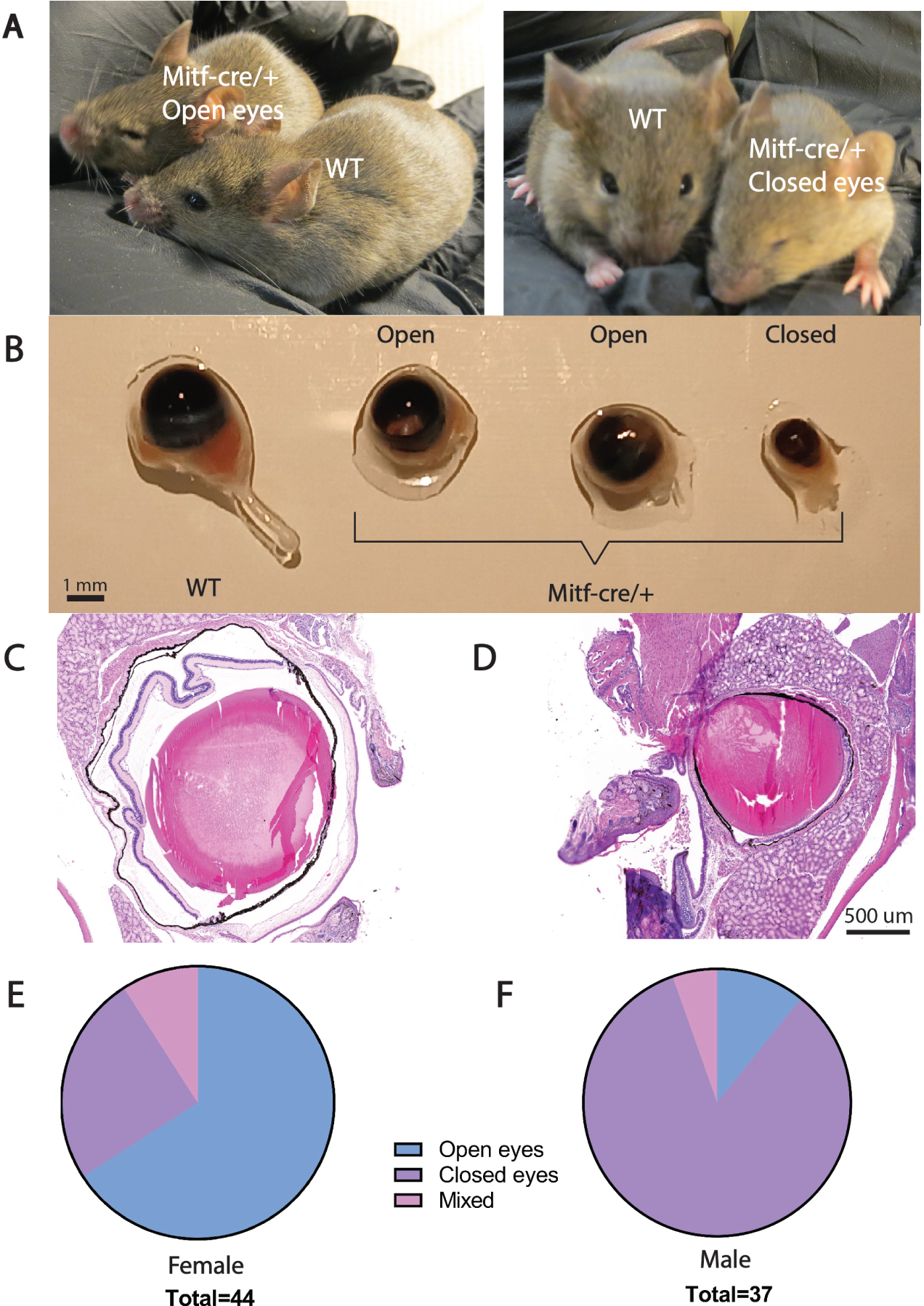
*Mitf-cre/*+ mice exhibit a range of ocular phenotypes. **(A)** Female *Mitf-cre/+* mice with small, but open eyes (left), or closed eyes (right), compared to wildtype (WT) littermates. **(B)** Eyes dissected from female WT and *Mitf-cre/+* mice (n=4) with open or closed eyes, as indicated. **(C)** Coronal H&E section of a *Mitf-cre/+* open eye within the intact skull. **(D)** Coronal H&E section of a *Mitf-cre/+* closed eye within the intact skull. **(E-F)** Pie charts showing the proportion *Mitf-cre/+* mice with open eyes, closed eyes, or a mixed phenotype (one eye open, one eye closed), based on the analysis of complete litters. Female mice are more likely to have open eyes.

Figure 2B illustrates the range of ocular sizes that occurred in the *Mitf-cre* colony, as seen in whole enucleated eyes. Although *Mitf-cre*/+ mice with completely closed eyelids suggested anophthalmia, there was usually a small sac-shaped eye that could be retrieved when dissecting from inside the skull (for example, see eye on the far right in **Figure 2B**). In addition, histological analysis of intact heads confirmed that most closed eyed *Mitf-cre* mice have a eye within the skull (**Figure 2C,D**). Interestingly, there was a sex difference in the severity of microphthalmia (**Figure 2E,F**). Male *Mitf-cre* mice had 16 times greater odds of having closed eyes compared to females (OR = 15.54, 95% CI [4.57, 52.77], p < 0.0001). We always cross female *Mitf-cre* mice with open eyes to C3HeB/FeJ males to maintain our colony, in case epigenetic effects are playing a role.

We also enucleated eyes from *Mitf-cre/+* female mice with open eyes and compared them to female wildtype eyes using transverse sections taken at the middle of the eye and stained with H&E (**Figure 3A-B**). We found that these *Mitf-cre/+* eyes were 22% smaller compared to wildtype (p=0.0045; Kolmogorov-smirnov test) (**Figure 3C**). The lens was often a relatively normal size. However, the outer nuclear layer of the retina was notably thinner (**Figure 3B**). Moreover, the pigmented layer of the eye (RPE and choroid) was 41% smaller (p=0.0002; Kolmogorov-smirnov test) (**Figure 3D**). Despite an apparent thickening of the ciliary body in the *Mitf-cre/+* eyes, the iris was 77% shorter when measured from the ciliary margin to the distal tips of each iris (p= 6.1 x 10^-7^; Kolmogorov-smirnov test) (**Figure 3E**).

**Figure 3.**
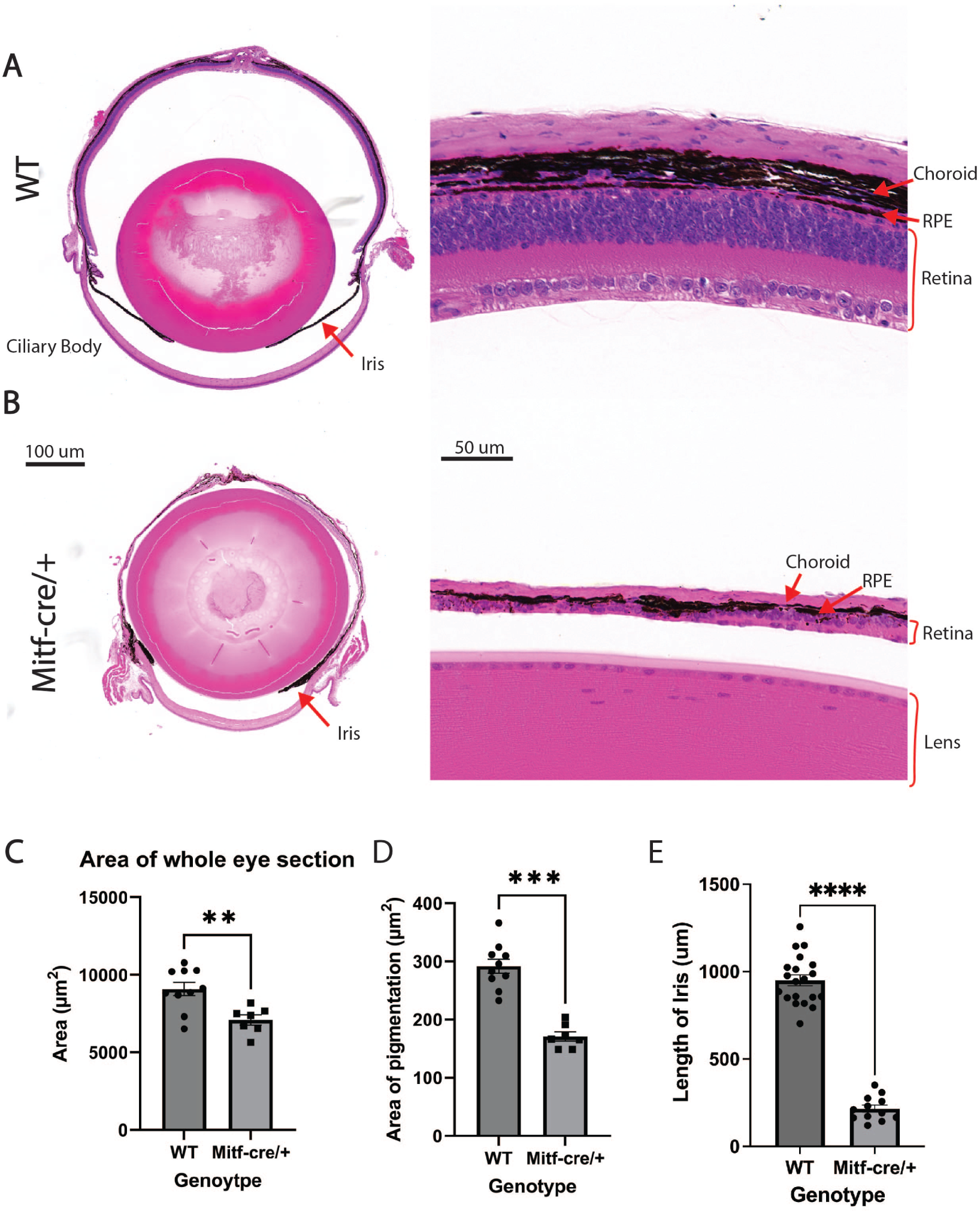
Female *Mitf-cre/+* mice with open eyes exhibit smaller eye size and thinner pigmented layers. **(A)** Transverse H&E stained section of a female WT eye. **(B)** Transverse H&E section of an open female *Mitf-cre/+* eye. **(C)** Comparison of the area of the whole eye in WT versus open *Mitf-cre/+* eyes. **(D)** Comparison of the area of the pigmented layer in WT versus open *Mitf-cre/+* eyes. **(E)** Comparison of the length of the iris in WT versus open *Mitf-cre/+* eyes.

To determine whether these phenotypic effects were apparent during early ocular development, we examined *Mitf-cre/+* and wildtype embryos at E13.5. Like adult mice, *Mitf-cre/+* embryos exhibited a range of eye sizes (**Figure 4**). A common finding in all whole mount *Mitf-cre/+* embryos was a notch of missing pigmentation on the ventral side of the eye, with an overgrown pigmented area on the dorsal side, consistent with coloboma (**Figure 4B-E**). Correspondingly, H&E stained cryosections of these eyes sometimes revealed a gap where pigmented layer was discontinuous (**Figure 4B,C,D**). Also, the pigmented tissue abnormally extended back towards the brain in a small line (see **Figure 4C,E**, whole mounts). In the most severely affected eyes, there was an absence of a lens (**Figure 4D,E**, sections) and the eye tissue that was there was not close to the surface ectoderm (**Figure 4E**). Lastly, there was unequal symmetry within eyes (**Figure 4B,C,D** sections).

**Figure 4.**
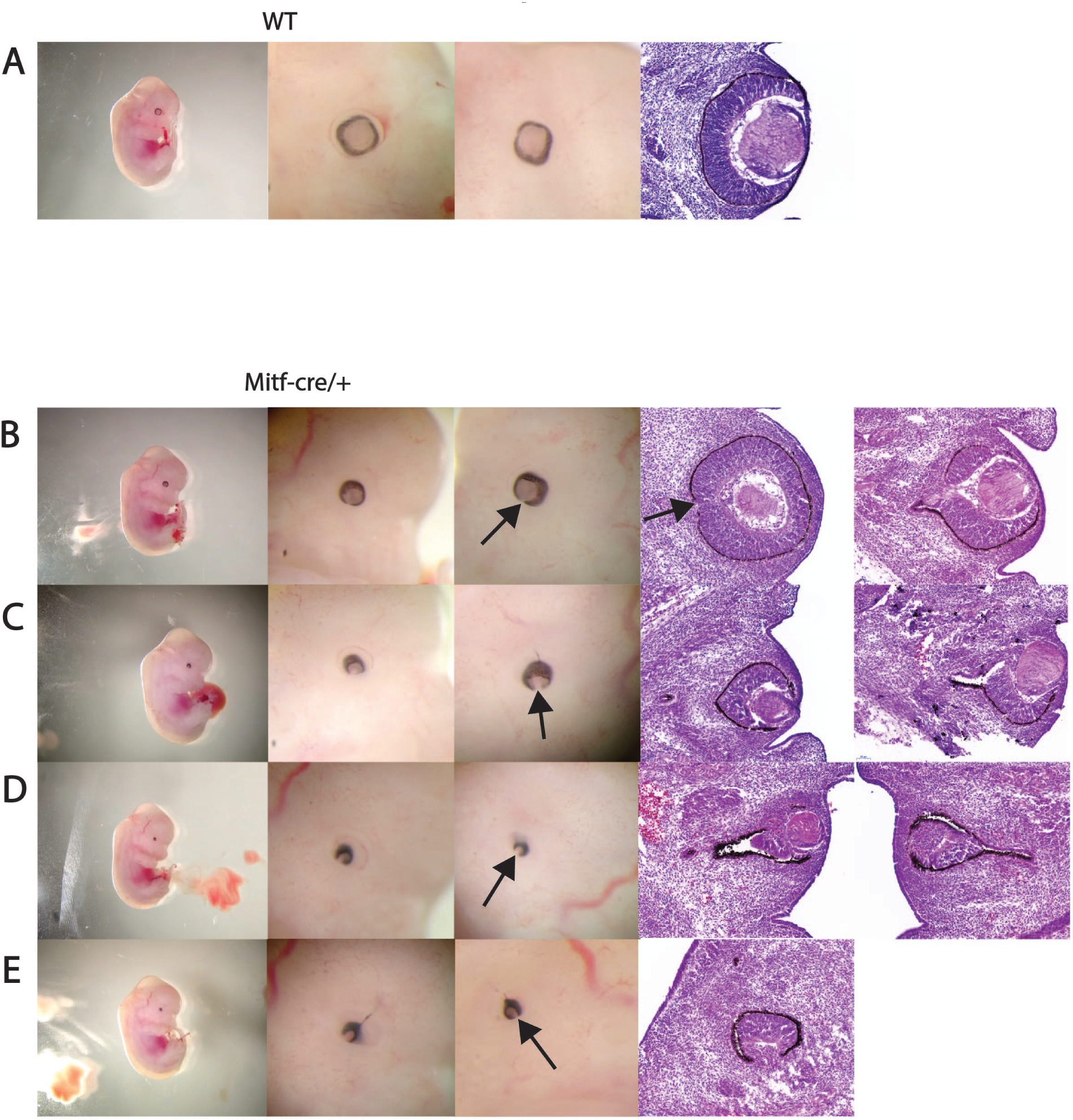
*Mitf-cre/+* E13.5 embryos display abnormal ocular development. **A-E)** Whole mount and H&E stained sections of a WT embryo (A) or *Mitf-cre/+* embryos (B-E) exhibiting a range of eye defects. On the left side, whole mounts are shown, along with the left and right sides of each eye in close up. Arrows indicate the gap in pigmentation on the ventral surface of the *Mitf-cre/+* eyes, consistent with coloboma. On the right side, H&E stained sections of eyes are shown.

To map the location of the *Mitf-cre* transgene insertion into the mouse genome, we submitted viable frozen splenocyte samples from *Mitf-cre* transgenic mice to Creative Bioarray for transgene mapping using targeted locus amplification (TLA) and next generation sequencing^30^. Transgene mapping and integration site characterization is possible through TLA by exploiting the principles of proximity ligation, which assumes that neighbouring sequences will be preferentially crosslinked and ligated together^30,31^. This method involves crosslinking, ligation and amplification of genomic DNA containing the fragment of interest or anchor fragment^30^. Primers specific to the transgene sequence are used as the anchor for PCR. The amplified products can then be sequenced and mapped to the genome or elements of the transgene to determine vector integrity, site of integration, and structural rearrangements^30,31^.

In our study, four independent primer sets were used to amplify anchor fragments: one specific for the Cre element and three specific for the mouse *Mitf* locus from chromosome 6 contained in BAC clone RP23-271N22 (**Table 1**). To determine the integrity of the transgene at its integration site, anchor fragments from the 4 primer sets were mapped to the mouse genome, the Cre sequence or the pBACe3-6 backbone.

**Table 1.**
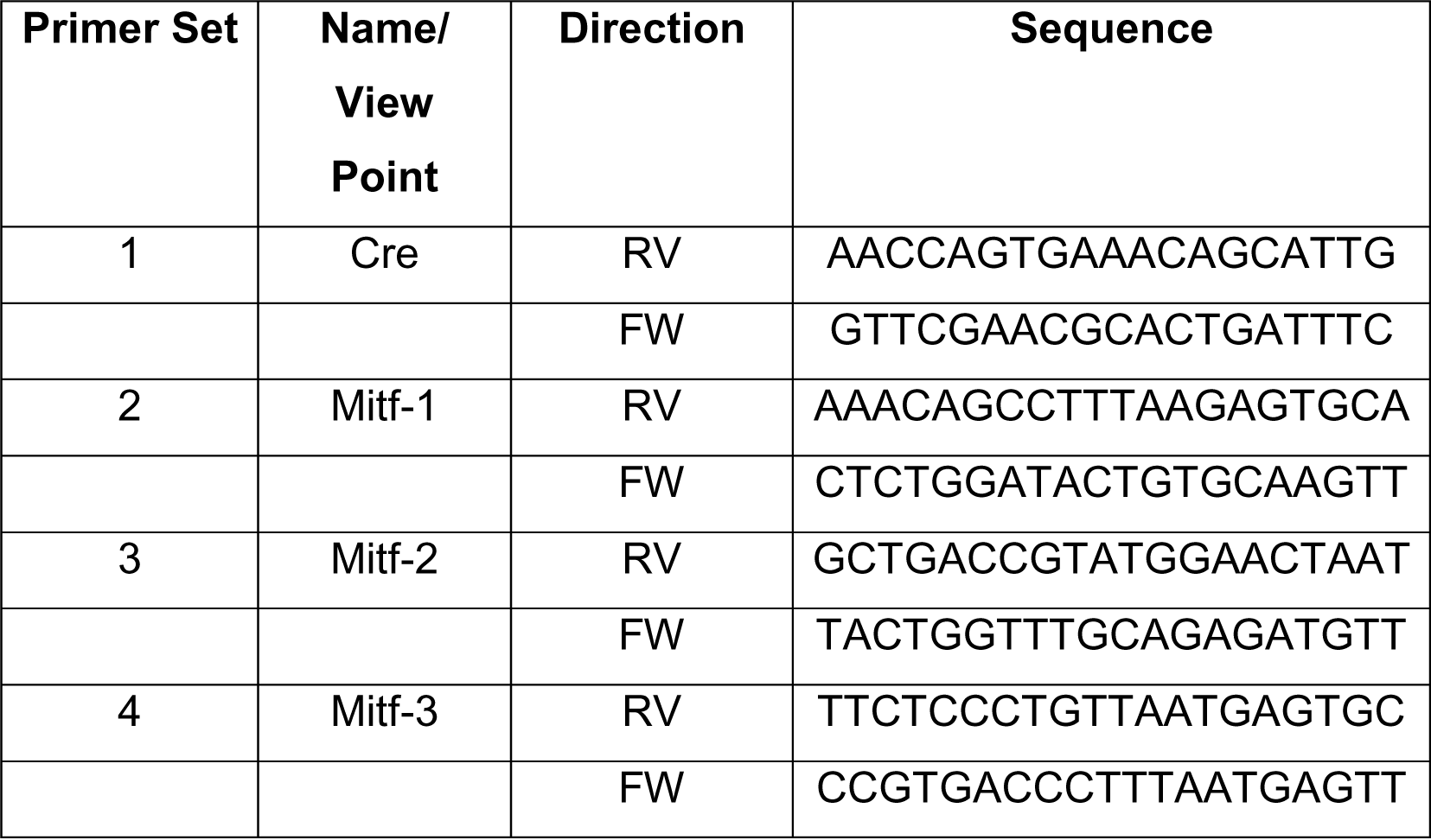
Anchor primer sequences.

Next generation sequencing (NGS) coverage was observed in the mouse genome from chr6: 97,882,418 - 98,067,774 [GRCm39] (**Figure 6C**), across the Cre sequence and across the backbone sequence, pBACe3-6. Therefore, 185.356 kb of chromosome 6, the Cre element, and the pBACe3 backbone were all successfully integrated. The coverage profile identified the BAC backbone connection to chromosome 6 at 98,067,774 [GRCm39], as well as the *Cre* gene insertion at the Mitf-M translation start site (chr6:97,968,884 [GRCm39]), validating the work done by Alizadeh *et al*^26^.

Based on the TLA data, BAC RP23-271N22 contained a larger portion of chromosome 6 than previously thought. To confirm this, we obtained the draft sequence of RP23- 271N22, which is now available from the NCBI (https://www.ncbi.nlm.nih.gov/nuccore/18092968) in 14 unordered pieces. Although there are gaps, we aligned most of the sequence to the mouse genome (**Figure 5A**). Those that matched chromosome 6 covered the region from Chr6:97,885,120 - 98,058,440 [GRCm39]), within the region identified by TLA. Inspection of this region on Ensembl shows that it includes annotations for the Mitf-H, Mitf-D and Mitf-B alternative promoters, not just Mitf-M (**Figure 5B)**. We also were able to directly align the Mitf-M, Mitf-H, and Mitf-B promoter sequences from Udono *et al.* (2000)^32^, and the Mitf-D promoter sequence from Takeda *et al.* (2002)^2^, to the RP23-271N22 draft sequence.

**Figure 5.**
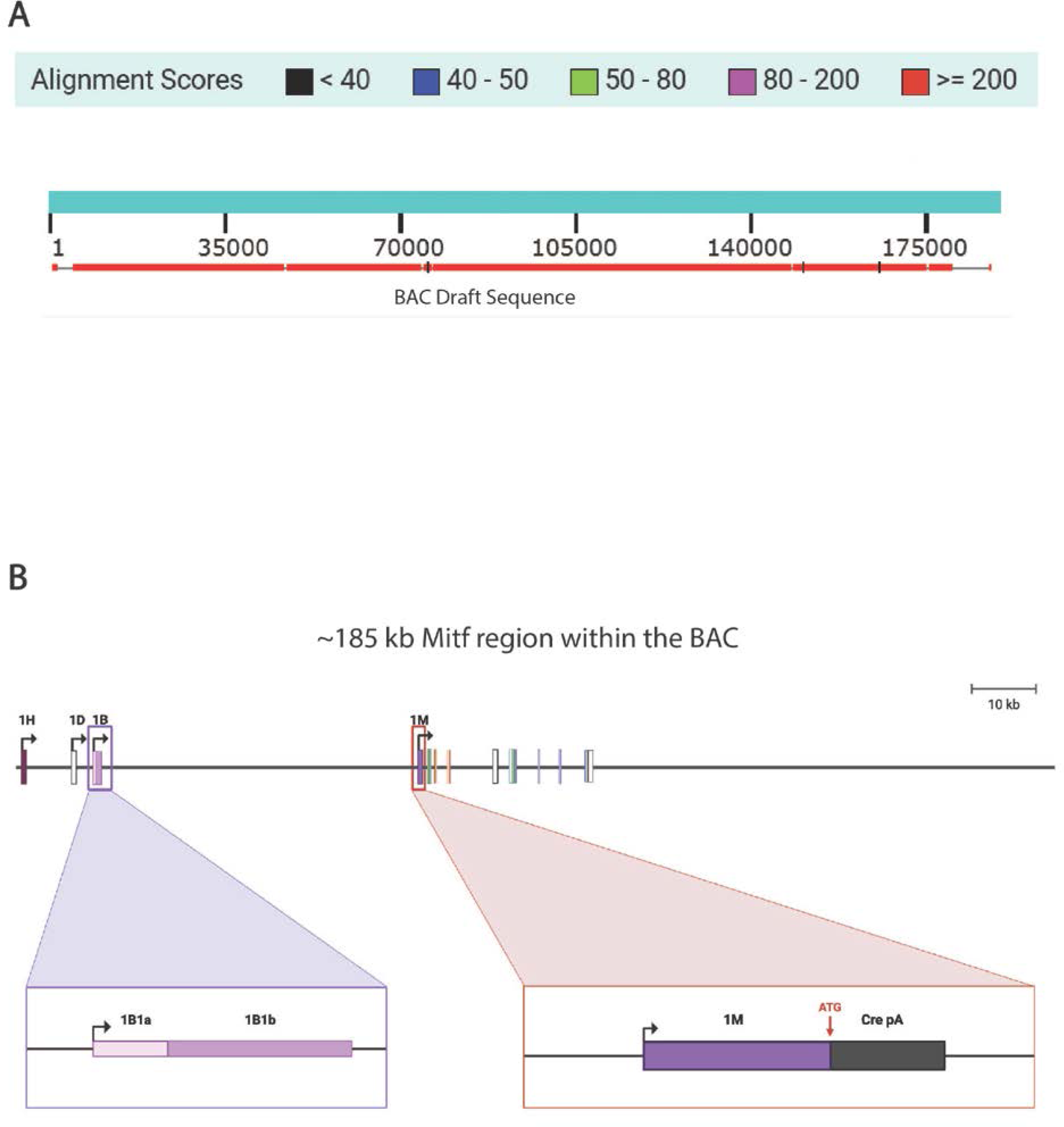
The *Mitf-cre* transgene carries three *Mitf* alternative promoters in addition to Mitf-M. **(A)** Graphic depicting the strength of alignment between the RP23- 271N22 draft sequence and mouse chr6:97,882,418-98,067,774 [GRCm39]. **(B)** Schematic showing the selected components of the *Mitf-cre* transgene. Mitf-H, Mitf-D, and Mitf-B alternative promoters are located upstream. Transcripts from the Mitf-H and Mitf-D promoters have been shown to splice to segment 1B1b, while transcripts from Mitf-B include it (shown in purple box). The Mitf-M promoter is downstream. In the engineered vector, Cre replaces the coding sequence of Mitf-M exon 1, at the ATG (shown in pink box).

**Figure 6.**
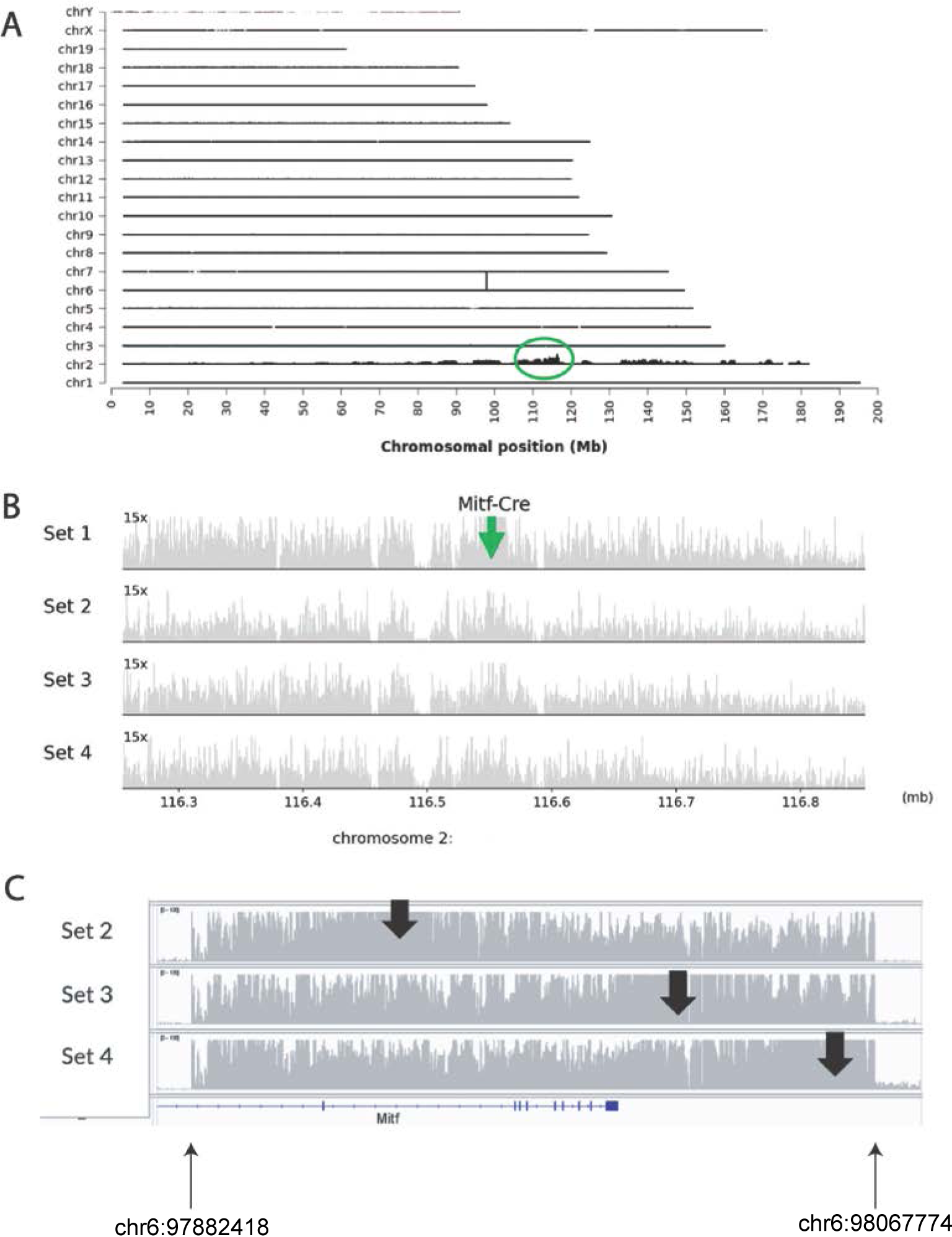
The *Mitf-cre* transgene inserted into an intergenic region on Chromosome 2. **(A)** Transgene mapping (locus amplification and sequencing) coverage across the mouse genome using primer set 1. Similar results were found with primer sets 2-4. The chromosomes are indicated on the y-axis and the chromosomal position on the x-axis. The identified integration site on chromosome 2 is circled in green. **(B)** Sequence coverage (in grey) across the BAC integration site on chromosome 2. The green arrow indicates the identified insertion site, detected in sequencing reads that link the BAC to the genomic sequence. The coverage profile shows that no genomic rearrangements have occurred in the region of the integration site. (**C**) NGS sequencing coverage (in grey) across mouse chromosome 6, indicating that there is 185 kb of chr6: 97,882,418 - 98,067,774 [GRCm39] on the BAC, RP23- 271N22. The *Mitf* gene is shown in blue beneath, the last exon is to the right.

Another important piece of information gained from TLA analysis was the integration site of the transgene into the mouse genome, which was identified to be chr2:116,384,742 [GRCm39] (**Figure 6A-B**). The copy number was estimated based on the coverage ratio between the BAC at chr6:98,067,774 and the genome at chr6:98,067,775 [GRCm39]. The coverage on the BAC-side is much higher than on the genome-side of the integration site (roughly 13 fold). The copies of the transgene integrated was thus estimated to be 13. Taken together, ∼2400 kb was inserted at chr2:116,384,742.

The integration site on chromosome 2 corresponds to an intergenic region (**Figure 7A**), thus no disruption of protein coding sequence occurred as a consequence of the *Mitf-cre* transgene insertion. Insertions within intergenic regions are historically presumed to have a negligible effect on endogenous gene expression^33^. However, work by Pal-Bhadra *et al*.^34^ has suggested that high copy number transgenes can instigate homology mediated co-suppression of its unlinked endogenous counterpart. In addition, other studies have proposed that transgenes can affect neighbouring genes in *cis* through a distance dependent manner due to competition for transcriptional machinery and transcription factors, as well as disruption of topologically associated domains (TADs)^31,33,35–37^.

**Figure 7.**
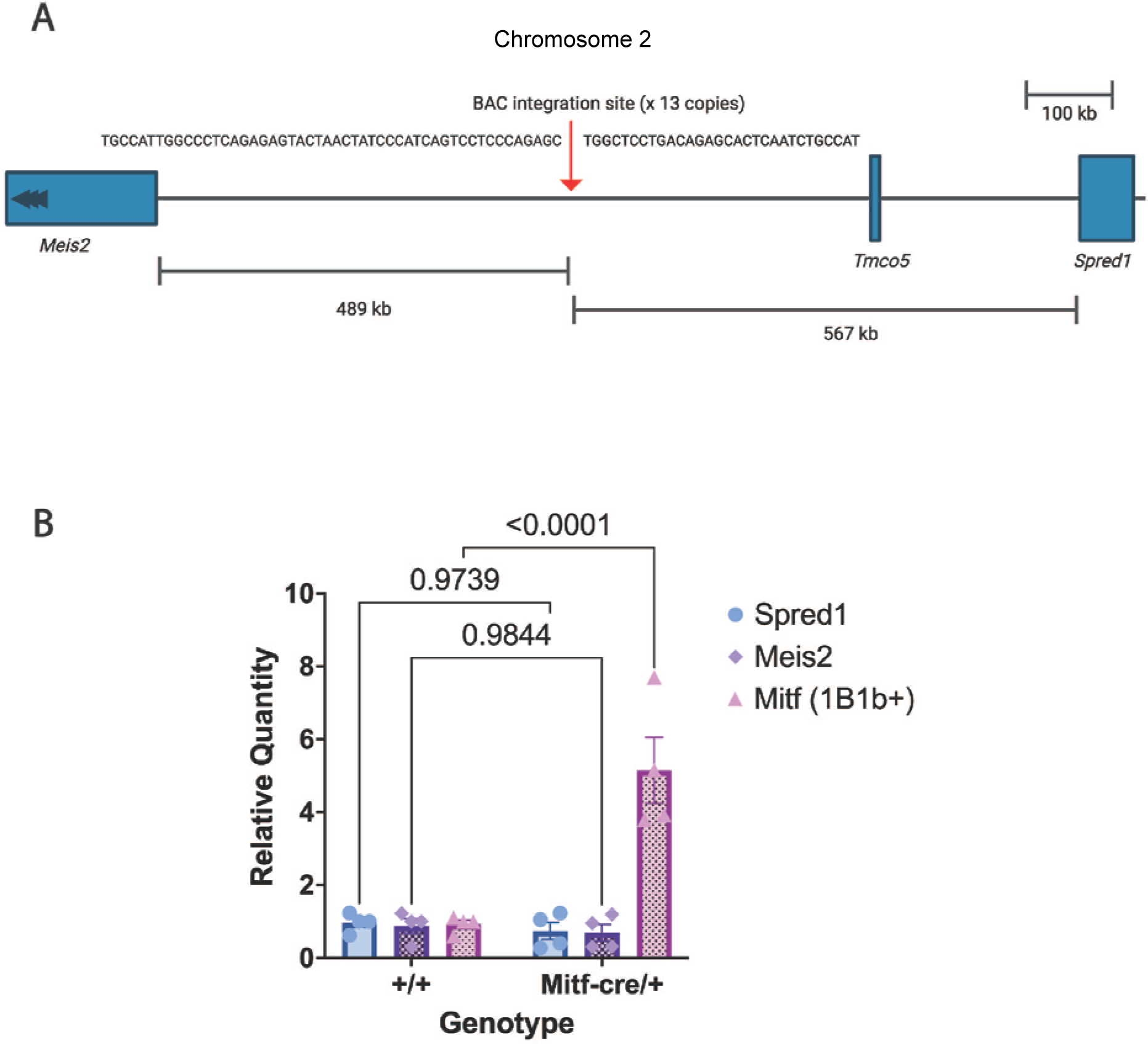
*Mitf* 1B1b containing transcripts are upregulated in *Mitf-cre/+* mouse eyes. **(A)** Schematic of the *Mitf-cre* transgene integration site on chromosome 2 and the relative position of the nearest genes. **(B)** RT-qPCR results comparing the relative quantity of *Spred1*, *Meis2*, and *Mitf* exon 1B1b containing transcripts in WT and *Mitf-cre/+* eyes, normalized to *Ppia*.

Using the Mouse Genome Informatics (MGI) database, we queried genes located on chromosome 2 that have been linked to microphthalmia in the literature (**Table 2**). We were interested to find that the transgene inserted in between two of them, *Meis2* and *Spred1* (**Figure 7A**). *Meis2* is detected in the optic cup and at the presumptive neuroepithelium of the ciliary margin, and throughout the retina^38^. Meis2 has been proposed to act as a transcription factor required for Vsx2 expression, promoting retinal progenitor cell specific gene expression while repressing ciliary margin specific genes^38^. Meis2 deficiency leads to the expansion of the ciliary margin at the expense of neural retina. Deletion of the Meis2 binding sites at *Vsx2* enhancers led to abolition of *Vsx2* expression and microphthalmia. In *Spred1* deficient mice, ocular defects have been reported with the most severe cases presenting with disruption of the optic cup and coloboma^39^. Spred1 and Spred2 work in concert during eye development. *Spred1/2* double KO’s present with a small lens (microphakia), which reportedly resolves with age^40^.

**Table 2.** Genes linked to microphthalmia on mouse chromosome 2.

| Gene Symbol | Transcript position on Chr. 2 [GRCm39] | Distance from Transgene (kb) | Citation |
| --- | --- | --- | --- |
| *Pax6* | 105,499,245-105,527,709 | 10,857 | Cunha 2019 |
| *Lgr4* | 109,747,992-109,844,602 | 6540 | Siwko 2013, Weng 2008 |
| *Meis2* | 115,693,545-115,896,320 | 489 |  |
| *Spred1* | 116,951,855-117,012,760 | 567 | Wazin 2020 |
| *Nphp1* | 127,582,652-127,630,817 | 11,198 | Matthias 2021 |
| *Stk35* | 129,642,437-129,674,207 | 13,258 | Miyamoto 2018 |
| *Jag1* | 136,923,376-136,958,564 | 20,539 | Anderson 2018; Tien 2012; Kiernan 2007 |
| *Csnk2a1* | 152,068,759-152,123,772 | 35,684 | Murakami 2021; Lou 2008 |
| *Tbc1d20* | 152,135,748-152,155,916 | 35,751 | Liegel 2013 |
| *Id1* | 152,578,171-152,579,330 | 36,193 | Du 2011 |
| *Asxl1* | 153,187,749-153,245,927 | 36,803 | Abdel-Wahab 2013 |

Together, this data suggests two possible causes of eye defects in *Mitf-cre/+* mice. First, the transgene exists within the same TAD as *Meis2* and/or *Spred1*, which increases competition for transcription factors and transcriptional machinery, leading to a reduction of *Meis2* and/or *Spred1* mRNA expression, and ultimately microphthalmia. Second, the BAC contains enough upstream regulatory sequences to enable expression of Mitf-H, Mitf-D, and/or Mitf-B from the transgene itself. The Cre recombinase sequence utilized to make this transgenic line contained a polyadenylation sequence to prevent expression of Mitf-M. This is located within Mitf-M exon 1, which is spliced over by the Mitf transcripts initiating upstream. Thus, the upstream Mitf-H, Mitf-D, and Mitf-B promoters may be able to express their full transcripts, splicing into the intact common exon 2 and remaining exons of *Mitf*, which are also on the transgene. Over-expression of the Mitf-H and/or Mitf-D isoforms might disrupt eye development.

To determine whether the phenotypic effects observed in the *Mitf-cre/+* mice are due to excessive expression of Mitf-H, Mitf-D, and/or Mitf-B or reduced expression of *Meis2* and/or *Spred1*, we performed RT-qPCR. We compared the relative expression of *Spred1*, *Meis2*, and *Mitf* 1B1b containing transcripts in *Mitf-cre/+* and wildtype (WT) whole mouse eyes. Note that transcripts from the Mitf-H and Mitf-D promoters splice to segment 1B1b, while transcripts from Mitf-B include it (**Figure 5B**). There were no differences in expression of *Spred1* or *Meis2* between *Mitf-cre/+* and WT mouse eyes. However, the results showed a 5-fold increase in *Mitf* 1B1b containing transcripts in *Mitf-cre/+* eyes compared to WT eyes (p<0.0001, two-way ANOVA) (**Figure 7B**). Therefore, we think that the mostly likely cause of the eye defects in *Mitf-cre* mice is an over-expression of the Mitf-H and/or Mitf-D isoforms in the eye.

## DISCUSSION

The *Mitf-cre* transgenic mouse line was engineered as a tool to study pigment cell biology using the melanocyte specific promoter, Mitf-M, to drive expression of Cre recombinase. While the expression of Cre was successfully melanocyte specific, severe microphthalmia was present in 100% of the progeny on the initial C57Bl/6 background, with eyelids closed. 5-10% of C57Bl/6 and related strains sporadically develop microphthalmia or anophthalmia, due to polygenic disease^41^. Previous work has shown that mice carrying mutations conferring microphthalmia exhibit decreased penetrance when the relative percentage of C3H genetic background increases^42^. Indeed, it was not until the *Mitf-cre* line was bred onto the C3HeB/FeJ that mice with open eyes and relatively normal sized eyes were observed. However, our observations here demonstrate that there are ocular abnormalities within even seemingly normal sized eyes. Therefore, the penetrance of microphthalmia on the C3HeB/FeJ genetic background may not have decreased, but the C3H genome has evidently modified the expressivity of the *Mitf-cre* transgene phenotype, as *Mitf-cre/+* mice present with a wider range of ocular phenotypes. We also determined that expressivity is modified by the sex of the animal.

We examined the effects of the *Mitf-cre* transgene on eye development in mice that are on the inbred C3HeB/FeJ background. We found that among the open *Mitf-cre/+* eyes, eye size was reduced 22% overall, with a reduced thickness of both the neural retina and RPE/choroid layer, and a shorter iris. A range of eye size was also observed at E13.5 in embryos. *Mitf-cre/+* eyes invariably showed a notch of missing pigmentation on the ventral side of the eye, with an overgrown pigmented area on the dorsal side, consistent with coloboma, or failure to close the optic fissure. Also, pigmented tissue can be seen to abnormally extend in a small line from the eye back towards the brain in some embryos. There was an unequal symmetry to the eyes and a failure of the most severely affected eyes to produce a lens, perhaps because the OV did not make it to the surface ectoderm to induce lens formation.

To determine the molecular cause of the eye phenotype in *Mitf-cre* mice, we used TLA and NGS and discovered that 13 copies of the *Mitf-cre* transgene inserted into an intergenic region of chromsome 2 at 116,384,742 [GRCm39]. Our analysis revealed the BAC to contain more alternative promoters of *Mitf* than originally thought. The presence of these additional *Mitf* promoters enables transcription of one or more of the *Mitf* isoforms containing exon 1B1b, namely, Mitf-H, Mitf-D, and Mitf-B. Through RT-qPCR, we demonstrated these additional promoters increase the expression of *Mitf* 1B1b containing transcripts in the *Mitf-cre* eye. Due to the expression boundaries that must be established and maintained throughout ocular development, the excess *Mitf* transcripts produced from the *Mitf-cre* transgene may be disrupting Mitf dosage requirements, ultimately giving rise to an abnormal phenotype (Figure 8 A-C). Two eye related genes closest to the transgene insertion site, *Meis2* and *Spred1*, did not change their expression.

**Figure 8.**
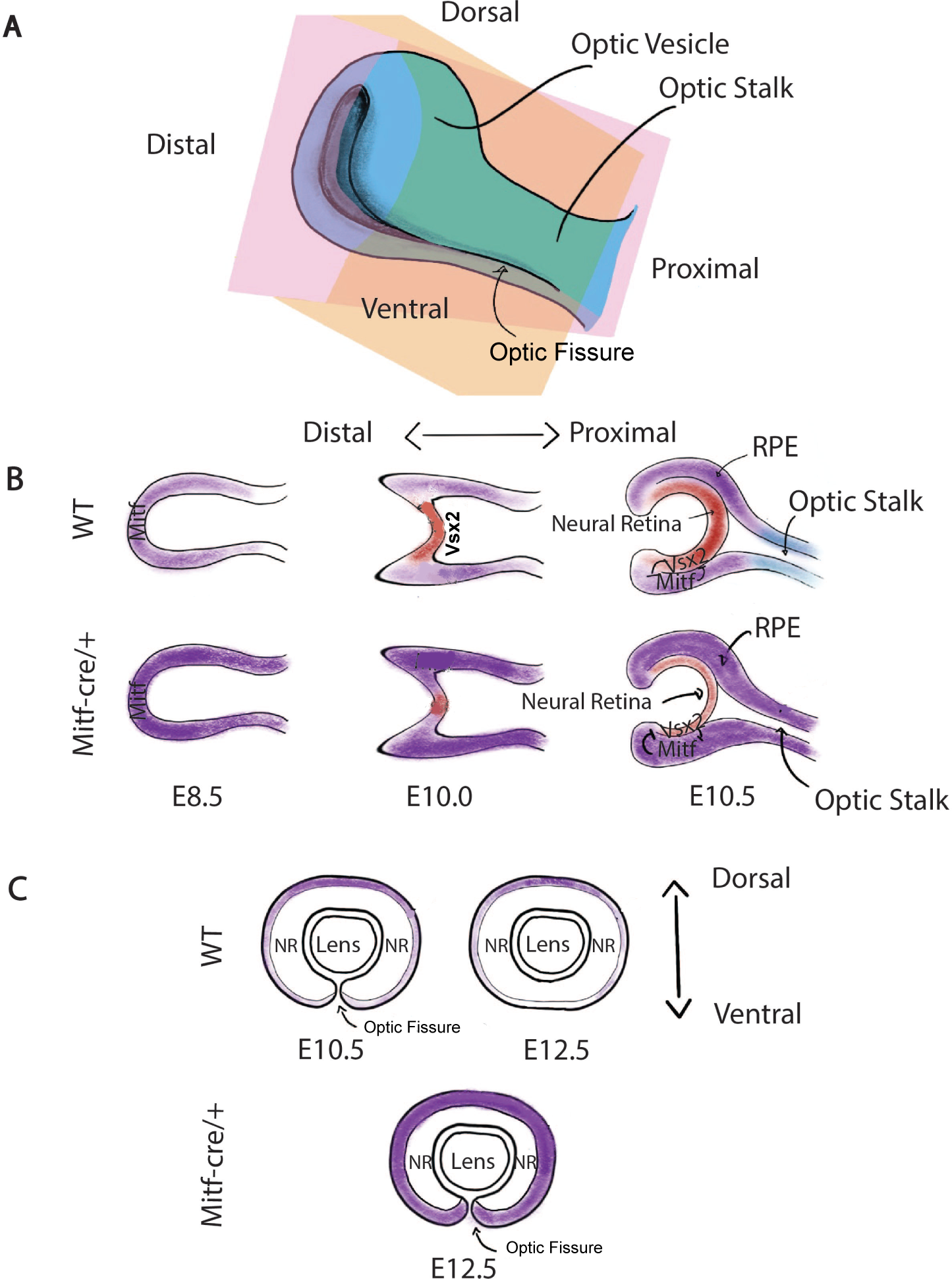
Model for *Mitf-cre* induced micropthalmia. **(A)** Schematic of mouse ocular development at E10.5 depicting the coronal and sagittal planes of reference as orange and pink respectively. **(B)** Comparing the effect of *Mitf-cre* on mouse ocular development at E8.5-10.5 as viewed from the sagittal plane. Mitf-H and Mitf-D isoforms play the most important role in retina and pigment epithelium development. The extra copies of the Mitf-H and Mitf-D isoforms on the *Mitf-cre* transgene may overwhelm the ability of Vsx2 to down-regulate *Mitf* properly. In addition, the over-expression of Mitf may inhibit *Vsx2* expression. The failure to exclude *Mitf* from the margins of the optic fissure leads to upregulated RPE fate and failed optic fissure closure. **(C)** Comparison of WT vs *Mitf-cre* ocular development at E10.5-12.5 as viewed from the coronal plane.

Multiple reports have suggested that the failure to down-regulate *Mitf* during ocular development can have detrimental effects on the developing eye. Prolonged expression of *Mitf* due to a mutation rendering its repressor, Vsx2, nonfunctional, leads to hypo-proliferation of the retina, microphthalmia, and the absence of bipolar interneurons in mice^14,20^. Moreover, neural retina progenitors exposed to higher ratios of Mitf:Vsx2 differentiated into an RPE like pigmented monolayer rather than neural retina, while still expressing neural retina markers^20^. By making use of the *Mitf ^mi-rw/mi-rw^* mouse mutant that deletes the Mitf-H, Mitf-D and Mitf-B promoter cluster, Bharti *et al.* also demonstrated that the Mitf-H and Mitf-D isoforms play the most important role in RPE development^20^. The extra copies of the Mitf-H and Mitf-D isoforms on the *Mitf-cre* transgene may overwhelm the ability of Vsx2 to down-regulate *Mitf* properly in the eye. In addition, the over-expression of *Mitf* in chicks inhibited Vsx2 expression, suggesting a reciprocal inhibition^43^.The pigmented line mentioned above in the E13.5 *Mitf-cre/+* embryos suggests that the excess Mitf generated from the *Mitf-cre* transgene enabled transdifferentiation of neural retina progenitors into RPE-like cells.

Furthermore, cross sections of adult *Mitf-cre/+* mouse eyes revealed a thinner retina compared to WT. This corroborates previous work in chick embryos, where the over-expression of Mitf significantly decreased the number of BrdU-positive neural retina cells^43^. Further experiments illustrated that *Mitf* over-expression led to ectopic pigmentation and *Pax6* inhibition in the presumptive neural retina^43^. In mice lacking *Pax6*, the neural retina cells fail to differentiate and proliferate^44^.

The *Mitf^Vit^* mouse mutant is a G-to-A transition that leads to an aspartate to asparagine substitution at amino acid 222 (p.D222N). It appears to retain DNA binding activity, but may impair protein-protein interactions. It causes microphthalmia in homozygotes. RT-qPCR and immunolabelling showed that there was a transient increase in Mitf expression in the dorsal RPE between E10.5-13.5 in *Mitf^Vit^*, followed by progressive retinal degeneration secondary to RPE abnormalities^45^. *Mitf^vit^* mice exhibited dorsal thickening of the RPE layer at E13 due to hyperproliferation which persisted until at least P6^46,47^. The degree of RPE abnormality appeared to be a function of hyper-proliferation, as the increase in cell number led to multilayering and misorientation of RPE cells, ultimately leading to progressive hypopigmentation^46^. In the *Mitf-cre/+* mice, we observed an up-regulation of *Mitf 1B1b* containing transcripts in adult mouse eyes, and this persistent up-regulation may be contributing to the RPE thinning and retinal degeneration that was observed. Taken together, *Mitf-cre* may directly contribute to neural retina differentiation failure, which may be made worse by an abnormal RPE, which is required to support retina function (Figure 8B).

Further studies are needed to assess the impact of the up-regulated expression of Mitf-H, Mitf-D, and/or Mitf-B on the ocular gene regulatory network throughout the course of ocular development. In addition, the timing and mechanism of ocular degeneration after E13.5 remains in question. Male *Mitf-cre*/+ mice were much more likely than females to develop severe microphthalmia (closed eyes) and the cause of this sex difference is unknown. Whether stochastic or epigenetic effects are at play in the mixed eye phenotype (where left and right eye phenotypes differ) is also a very interesting avenue of research that could be explored in the mice. The findings here clarify and support the usage of *Mitf-cre* in driving oncogene expression and knockouts in melanocytes. The specific over-expression of the Mitf-H and Mitf-D isoforms, which are preferentially expressed in the RPE, presents a unique resource for those interested in eye development and coloboma.

## METHODS

### Mouse Husbandry and genotyping

The research described in this article was conducted under the approval of the UBC Animal Care Committee (UBC animal care protocol numbers A19-0152 and A23-0152, C.V.R). *Tg(Mitf-cre)7114Gsb* ("*Mitf-cre"*) mice have been maintained on the C3HeB/FeJ genetic background for >10 years. For genotyping, DNA from ear notches was isolated using the DNeasy Blood and Tissue kit (Qiagen) and amplified using PCR with HotStar Taq (Qiagen). The *Cre* genotyping primers are:

L:GCGGTCTGGCAGTAAAAACTA

R:GTGAAACAGCATTGCTGTCAC.

### Histology in adult tissue

Adult mouse eyes were fixed in Davidson’s fixative for 3 hours, followed by 10% formalin for 1 hour. Whole heads were fixed in 10% formalin overnight and were decalcified with EDTA. All samples were dehydrated, cleared, embedded in paraffin and sectioned at 5 μm, before staining using standard H&E techniques by WaxIt Histology services (Vancouver BC). Images were collected using an Axio Scan.Z1 slide scanner (Zeiss).

### OCT Sample Preparation of embryos

Optimal cutting temperature (O.C.T.)-embedded E13 embryos were used to analyze the developmental eye phenotype. Embryos were fixed in 10% formalin for 7 hours at 4°C in the dark, then washed in 1x Phosphate Buffered Saline (PBS) for 1 hour. Samples were then transferred to a 10% sucrose solution (in 1x PBS) for 1 hour, followed by a 30% sucrose solution (in 1x PBS) for 11 hours. The embryos were embedded head down in Scigen Tissue-Plus™ O.C.T. compound (Fisher Scientific) to allow for 10 μm transverse sections to be acquired using the Leica CM 1950 Cryostat. Once the eyes were reached, every second section was collected and mounted on Superfrost™ Plus glass microscope slides (Fisher Scientific).

### H&E Staining of embryos

The O.C.T sections of embryo eyes were stained with Hematoxylin and Eosin (H&E) to aid in visualization of cell nuclei and cytoplasm. Slides were first washed in 1x PBS for 30 minutes then rinsed twice in water for 2 minutes. The water was poured off and 400 μL of hematoxylin (BioLynx Inc.) was added to each slide and left for 1 minute. Slides were immediately rinsed with tap water. Slides were differentiated in an acid rinse solution (2% glacial acetic acid) and rinsed in tap water. An ammonium hydroxide-based blueing solution was then added to the slides followed by a rinse in tap water. Slides were then dehydrated in 10% increasing increments of ethanol from 30-80%, for 2 minutes at each concentration. Eosin (prepared in 80% ethanol and glacial acetic acid) was added to the slides for 10 seconds. Slides were immediately taken through 80% and 95% ethanol for 2 minutes each, followed by 5 minutes in 100% ethanol twice. Slides were then washed twice for 10 minutes with the clearing agent, xylene, and mounted in Eukitt mounting solution (Sigma). Images were collected using an Axio Scan.Z1 slide scanner (Zeiss).

### Transgene mapping

To map the location of the *Mitf-cre* insertion into the mouse genome, we submitted viable frozen splenocyte samples of *Mitf-cre* transgenic mice to Creative Bioarray for transgene mapping which utilizes targeted locus amplification (TLA) and targeted sequencing^30^. Splenocytes were isolated from 7-week-old *Mitf-cre/+* mice by collecting the whole spleen and forcing it through a 40 μm mesh. Cells were washed, centrifuged, and resuspended in red blood cell lysis solution. Then they were washed and centrifuged again, and resuspended in freezing solution with fetal calf serum and DMSO. Samples were frozen and stored at −80°C or and on dry ice during shipping.

Transgene mapping and integration site characterization was done with proximity ligation, which crosslinks and ligates neighboring sequences together^30,31^. This is followed by amplification of genomic DNA containing the fragment of interest, the anchor fragment^30^. By using primers specific to the transgene sequence as an anchor, TLA enables targeted amplification of the transgene and neighbouring sequences. The amplified products can then be sequenced using next generation sequencing technology and mapped to the genome and known elements of the transgene to determine vector integrity, site of integration, and structural rearrangements^30,31^.

### RNA isolation

For each mouse, one eye was enucleated with curved forceps and then placed on a petri dish sitting on ice while the lens was removed with fine forceps. The remainder of the eye was placed in cold 300 μL Trizol in a 1.5 mL eppendorf tube and disrupted for 5 minutes with a handheld homogenizer with an RNA-free disposable pestle. 700 μL more Trizol was added, the tube was vortexed and then held on ice while the second eye was disrupted as above. Then, both tubes were incubated for 5 minutes at room temperature. 200 μL of choloform was added to each, vortexed and incubated 3 minutes. At 4°C, the samples were spun at 25,000g for 15 minutes. The upper phases were placed into new tubes and precipitated with 500 μL ice cold isopropanol on ice for 10 minutes, followed by centrifugation at 25,000g for 10 minutes. The resulting pellets were washed with 75% ethanol, dried and resuspended in 20 μL of water. RNA quality and quantity was determined by an Agilent 2100 bioanalyzer. All RIN scores were above 9.2.

### RT-qPCR

RT-qPCR was performed using the Verso 1-step RT-qPCR Kit, SYBR Green, low ROX (Thermo Fisher, AB4106A) and a QuantStudio3 real time PCR machine (Applied Biosystems). Primers were designed to span exon boundaries using Primer3plus. Primers were first tested in standard curve analysis using RNA extracted from a normal eye to establish that each generated efficient amplification. RNA samples were extracted as above from 2 female WT mice (4 eyes) and 2 female open eyed *Mitf-cre/+* mice (4 eyes) at 8 weeks of age. 50 ng of RNA was used for each reaction. Amplicons from genes of interest were amplified using the following primers, in triplicate, alongside no-template and no RT enzyme controls. The expression of each gene of interest was normalized to ubiquitously expressed, *Hprt*.

### Meis2

Left primer in exon 9, right primer in exon 11

Meis2-L: CACGGGACTGACAATTCTGC

Meis2-R: GATCCCCATGTGTTGCTGAC

### Mitf 1B1b

Left primer in exon 1B1b, right primer in exon 2

Mitf 1B-L: CACCAGCCATAAACGTCAGC

Mitf 1B-R: GACATGGTGAGCTCAGGACT

### Spred1

Left primer in exon 5, right primer in exon 7

Spred1-L: TTGCAAAGCCAAGTCAGCC

Spred1-R: TCCGGATGTCTGTAGTCTGC

### Hprt

Left primer in exon 6, right primer in exon 7

Hprt-L: TTCCCTGGTTAAGCAGTACAGC

Hprt-R: TCTGGCCTGTATCCAACACTTC

### Statistical analysis

Data was analysed using the indicated tests with GraphPad Prism 8.4.3. Error bars represent the standard error of the mean.

## ACKNOWLEDGEMENTS

Funding for this research was provided by a grant from the Canadian Institutes of Health Research (CIHR), PJT-178178, to C.D. Van Raamsdonk. We thank Greg Barsh of Stanford University for the *Mitf-cre* mice. Figures created in part with BioRender.com.

